# SOFIA: an R package for enhancing genetic visualization with Circos

**DOI:** 10.1101/088377

**Authors:** Luis Diaz-Garcia, Giovanny Covarrubias-Pazaran, Brandon Schlautman, Juan Zalapa

## Abstract

Visualization of data from any stage of genetic and genomic research is one of the most useful approaches for detecting potential errors, ensuring accuracy and reproducibility, and presentation of the resulting data. Currently software such as Circos, ClicO FS, and RCircos, among others, provide tools for plotting a variety of genetic data types in a concise manner for data exploration and presentation. However, each of the programs have one or more disadvantages that limit their usability in data exploration or construction of publication quality figures, such as inflexibility in formatting and configuration, reduced image quality, lack of potential for automation, or requirements of high-level computational expertise. Therefore, we developed the R package SOFIA, which leverages the capabilities of Circos by manipulating data, preparing configuration files, and running the Perl-native Circos directly from the R environment with minimal user intervention. The advantages of integrating both R and Circos into SOFIA are numerous. R is a very powerful, mid-level programming language widely used among the genetic and genomic research community, while Circos has proven to be a novel software for arranging genomic data to create aesthetical publication quality circular figures. Producing Circos figures in R with SOFIA is simple, requires minimal coding experience, even for complex figures that incorporate high-dimensional genetic information, and allows simultaneous analysis and visual exploration of genomic and genetic data in a single programming environment.

## Introduction

Visualization is one of the best strategies for exploring, analyzing, and presenting data in common genetic and genomic studies such as linkage mapping, quantitative trait loci (QTL) mapping, association studies, and comparative genomics. These types of genetic analyses, especially those related to genetic mapping, generally involve a series of methodological steps such as creation of mapping populations, defining the type and number of markers to use, data cleaning, estimation of recombination, linkage group ordering and alignment, phenotyping, and the evaluation of the genotype-phenotype associations. Each step contains its own potential sources of error, and data visualization is an important means for detecting the introduced error and ensuring the resulting accuracy and reproducibility of the study.

Most packages and/or software for genetic analysis possess visualization tools for exploring general aspects of the data. For example, the commercial software, JoinMap (Stam 1993), which is widely used for genetic map construction, can display constructed linkage groups, alignments of linkage groups to compare marker position and linkage group structure between populations or species, and colorized phased genotypic data to facilitate exploration of recombination events and to detect genotyping errors in the mapping population. Other software for performing QTL and genome-wide association studies (GWAS), such as MapQTL (Van Ooijen 2009), r/qtl (Broman et al. 2003), and sommer (Covarrubias‐Pazaran 2016), provide functions for plotting LOD scores and *p*-values for detecting genotype-phenotype associations. However, integrating and arranging data from these genetic software and independent genetic analyses into single aesthetical images for simultaneous visualization remains challenging.

Circos (Krzywinski et al. 2009) is a novel software that addresses the challenges in visualizing genetic data by creating circular ideograms composed of “tracks” of heatmaps, scatter plots, line plots, histograms, links between common markers, glyphs, text, and etc. The flexibility of the software makes it suitable for rapid deployment in linkage and QTL mapping analyses and is especially useful for comparing genetic data between individuals, populations, and species. Circos, an open-source tool, has proven to be one of the most effective ways to display high-dimensional data, and it is one of the most used software (i.e. more than 2200 citations) for visualization in genetic and genomic research. However, the Perl-native Circos operates through a command-line interface that, while highly flexible, requires the user to have advanced computational skills. For users with little programing experience, this remains an obstacle to the routine implementation of Circos for data exploration and figure development.

In this paper, we present an R package, SOFIA, which is a powerful tool for visualizing genetic data that combines the advantages of the R programming language and Circos. Our package provides a pipeline for producing high quality images with the potential to integrate high-dimensional genetic data in an aesthetical and highly useful manner. Most importantly, unlike other available software, SOFIA is a tool that exploits most of the capabilities of Circos, but integrates it within the friendlier R programming language to allow simultaneous data analysis and visualization in a single programming environment.

## Software similar to SOFIA

To facilitate the use of Circos, additional tools and software packages have recently been created, such as Circoletto (Darzentas 2010), ClicO FS (Cheong et al. 2015) and RCircos (Zhang et al. 2013), which aim to provide a more user-friendly interface for using Circos. Unfortunately, each of these tools lacks some of the qualities and essential attributes of native Circos such as the flexibility, automatization, resolution, and robustness. For example, because running Circos from the Command Line can be challenging for computationally inexperienced users, tools like ClicO FS (Cheong et al. 2015) and Circoletto (Darzentas 2010) use a graphical interface (GUI) to facilitate the generation of Circos-like figures. While is true that the GUIs allow the easy formatting and structuration of the data for plotting Circos-like figures, these particular software remain too inflexible in terms of automatization, scalability, and personalization necessary for data exploration during preliminary analysis and final figure preparation. In fact, plotting recombination blocks (as heatmaps) for linkage groups in a genetic map with 1000 markers and 100 plants, a common preliminary analysis to identify genotyping errors leading to potentially false double-recombination events, could take a considerable amount of time if a GUI is used (plus the amount of time required for preparing all the numeric data files). The other available software, RCircos (Zhang et al. 2013), runs completely within the R programming language and produces Circos‐like figures in a more semi-automatized fashion, but it is very inflexible and lacks the formatting and configuration capabilities and resulting figures tend to be of lower quality compared to those produced by Circos (Figure 1).

**Figure 1.**
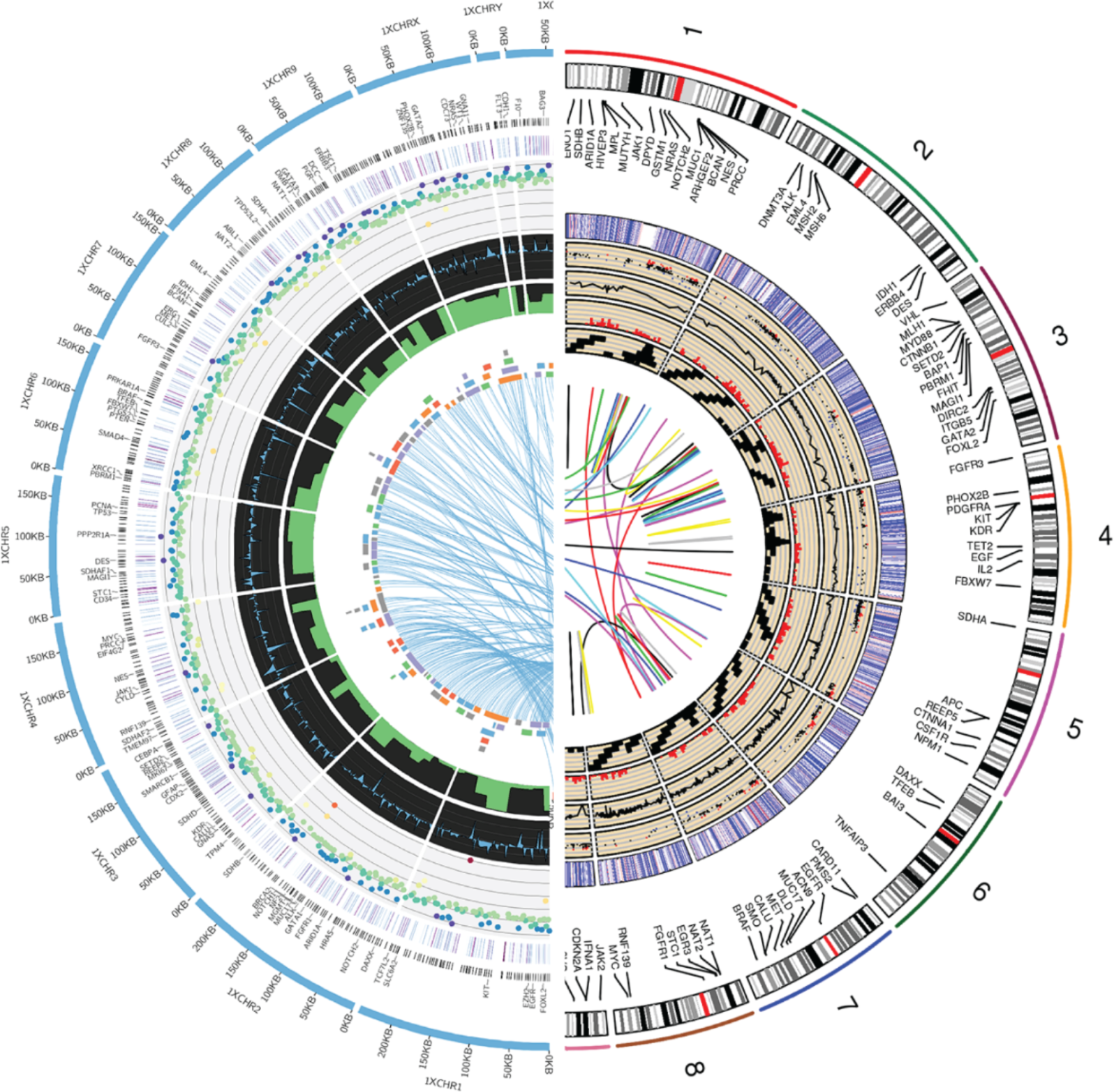
A comparison between a figure produced by Circos (through SOFIA, on the left) and RCircos (in the right). Similar data was used to generate both figures.

## Features and functionality of SOFIA

SOFIA is an R package that prepares the numeric data (i.e. 2D tracks including formatting) and configuration files (i.e. circos.conf) and then generates circular ideograms by running Perl-native Circos. SOFIA can be used by both experienced and inexperienced native-Circos by automatically preparing all necessary configuration files and then runing Perl-based Circos directly from R. Most of the functionality of Circos remains available in SOFIA, including all 2D tracks (scatter plots, line plots, glyphs, text, links, histograms and heatmaps), formatting, figure configuration, and etc. By running Perl native-Circos through R, SOFIA produces high-quality Circos figures while enormously reducing the required code (compared with other Circos-related software) and keeping most of the Circos functionality relating to formatting and flexibility. SOFIA provides a mid-level platform for easily generating high-quality Circos figures while maintaining the capabilities of R (Table 1).

**Table 1.** Comparison of general functionalities of available Circos-like software.

| Properties | Software |  |  |  |  |
| --- | --- | --- | --- | --- | --- |
|  | Circos | Circoletto | RCircos | ClicO FS | SOFIA |
| Author | Krzywinski M, 2009 | Darzentas, 2010 | Zhang et al., 2013 | Wei-Hien et al., 2015 | - |
| Platform | Perl | Online and Perl | RCircos | Online (uses native-Circos) | R (uses native-Circos) |
| Objective | Visualization of data in a circular fashion | Sequence similarity visualization | Visualization of data in a Circos-like fashion | Visualization of data in a circular fashion (produce Circos figures) |  |
| Data plotting capabilities | Supports scatter plots, line, histogram, heatmap, tile, connectors, links, and text labels. | Links and histograms | Supports scatter, line, histogram, heatmap, tile, connectors, links, and text labels |  |  |
| Image capabilities | High capabilities for formatting and ideogram configuration. | Built-in BLAST alignments. No other configuration functionality | Does not provide functionality for data processing and file preparation for running the package | High capabilities for formatting and ideogram configuration. |  |
| Automatization capabilities | Yes | No | Yes | No | Yes |

In the Circos-native version, automatic track plotting is supported, which facilitates the rapid configuration of data sets from multiple genetic analyses. In SOFIA, this feature can be implemented by simply iterating in R (with for loops, example) the data to be plotted as well as the location and formatting of the tracks in the circular ideogram. An important application of this approach is exploited, for example, for constructing representations of genetic linkage blocks (by using heatmaps).

Since very few code lines are required to produce SOFIA figures, as expected, some of the Circos functionalities (mainly those related with formatting) are not available in our package. However, running SOFIA produces all configuration files that can be further modified manually (through a text editor) to add other parameters not currently supported in SOFIA. When working with several SOFIA figures, it is very straightforward, in terms of organization, to keep a single code file (with very few code lines) to generate every figure without other manual user intervention.

To take advantage of the help resources and tutorials available for operating Circos (http://circos.ca/documentation/tutorials), we kept as much of the Circos syntax and logical flow in SOFIA as possible, especially in regards to formatting (for example color schemas). In our website (https://cggl.horticulture.wisc.edu/software/), we provide a library of figure templates that we have found to be very useful for exploring and analyzing data from linkage mapping, association studies, or any genomic study. We also provide samples of the figures produced by SOFIA as well as the R code for generating them (Figures 2 and 3; Supplementary File 1).

**Figure 2.**
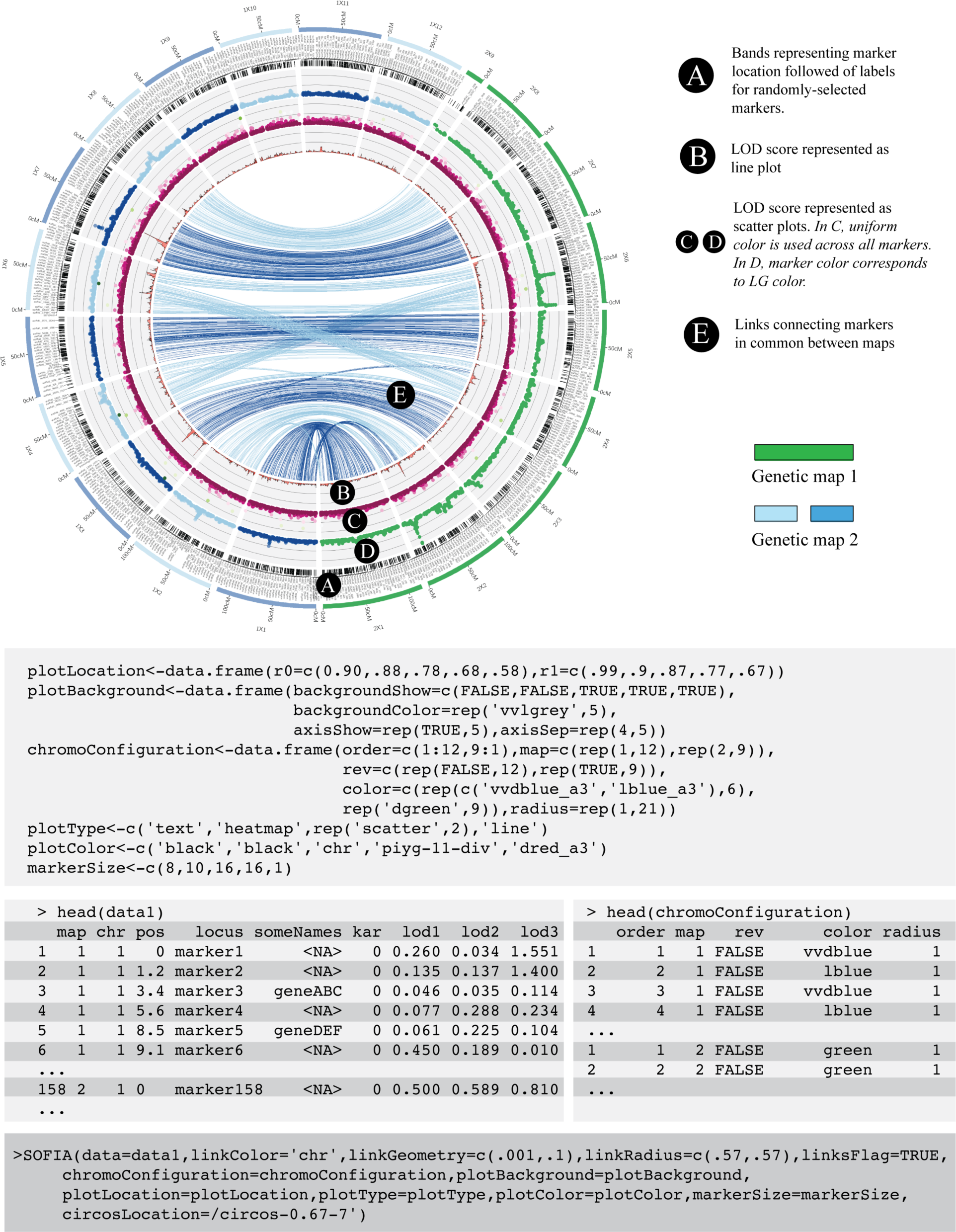
A Circos figure generated through SOFIA for visually comparing two genetic maps and locations of marker-trait associations identified through GWAS. All code required for producing the figure is presented in the gray boxes. First, the argument ‘plotLocation’ specifies the location of the 2D tracks within the figure (from 0 to 1, where 0 is the center of the image and 1 is where the linkage groups (LGs) are located). Then, ‘plotBackground’ controls the background characteristics such as color and separation between y-axis lines. ‘chromoConfiguration’ specifies the order, orientation and color of the LGs among the maps. The arguments ‘plotType’, ‘plotColor’ and ‘markerSizes’ control the properties of the plots for each of the 2D tracks to be plotted. Both the map and relevant data (i.e. LOD scores) must be merged and formatted as a single dataframe (as shown in the figure) which is the main input argument for running SOFIA. The R script for generating this figure can be found in Supplementary File 1.

**Figure 3.**
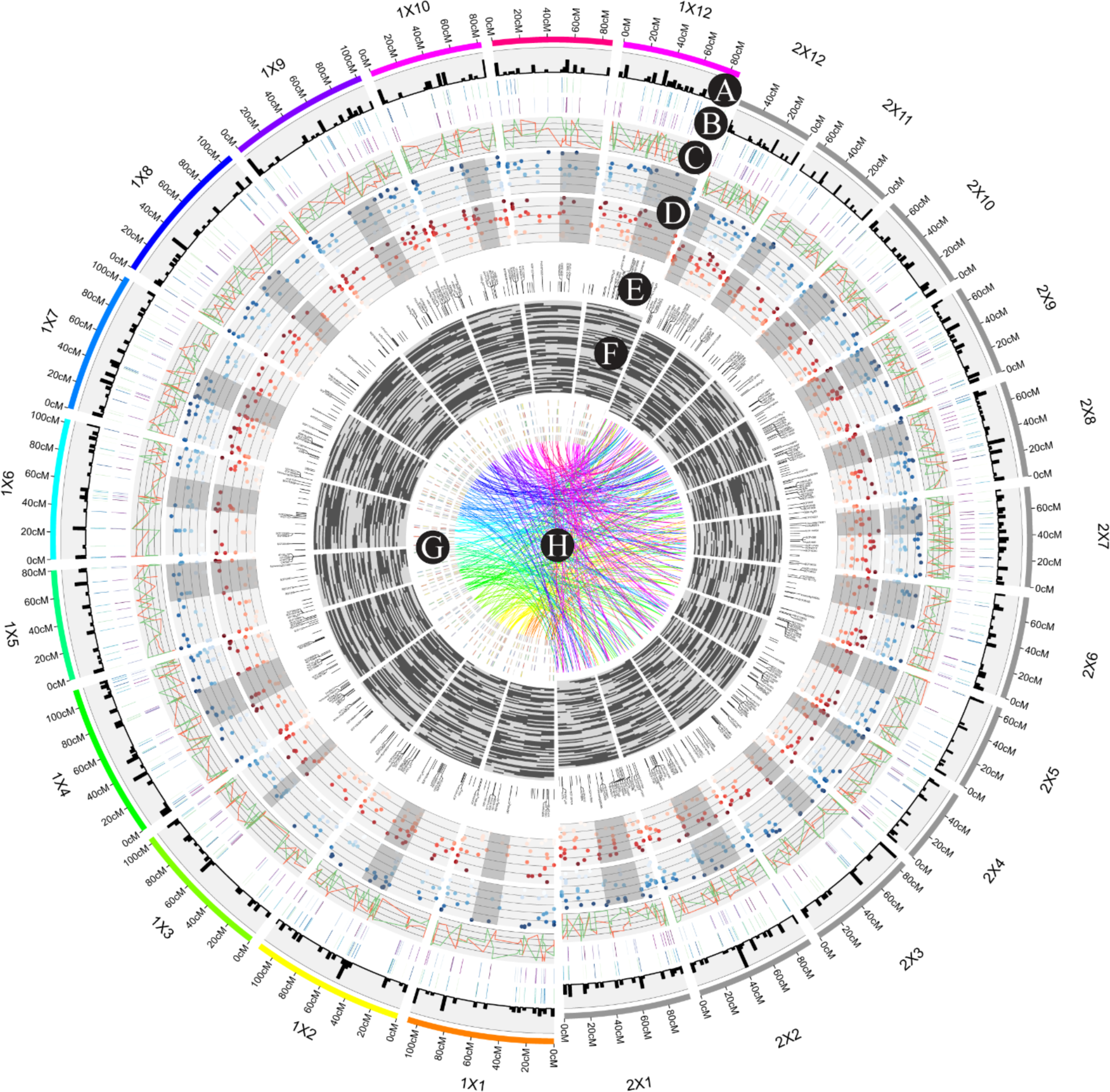
A Circos figure generated through SOFIA using a simple R script. Across rings, different types of genetic data are shown for two genetic maps comprised of 12 linkage groups each. In (A), the marker density is displayed as histograms, followed by (B) two heatmaps representing random data. Then, (C) a dataset plotted as lines in different colors and D) two scatter plots in which the centromeric region is highlighted. In the inner part of the figure, (E) text labels for some markers are displayed, followed by (F) recombination blocks for 74 genotypes. In (G), a set of tiles in different colors represents the genes across the linkage groups. In the inner part, (H) links connecting similar markers between maps are displayed. The R script for generating this figure can be found in Supplementary File 1.

In Figure 2, we show a typical example of how comparisons of marker-trait associations between genetic maps for two species or populations (with 12 and 9 LGs each) can be visualized with SOFIA. On each map, we display the presence/absence of genetic markers (black lines) as well as labels for randomly-selected positions (Figure 2A). Additionally, three plots (two scatter plots and a line plot) display log-transformed *p-values* scores from a GWAS study across all the linkage groups (Figure 2B, C, and D) followed by links connecting common markers between genetic maps and colored according to the linkage group in one of the maps (Figure 2E).

As we previously mentioned, SOFIA offers multiple options for plotting and formatting data in many different ways. Figure 3 serves as an example of a more complex figure that could be generated for publication purposes. In this figure, we show different types of data for two genetic maps; in the outer ring, a histogram showing the marker density is displayed (Figure 3A), followed by two heatmaps (in blue and purple) with random data (Figure 3B). Subsequently, a ring with datasets plotted as lines in different colors (Figure 3C) and two scatter plots in which the centromeric region is highlighted in darker grey color (Figure 3D). In the inner part of the figure, labels for randomly-selected markers are displayed (Figure 3E), followed by recombination blocks for 74 genotypes (Figure 3F). The “recombination blocks” plot type is one the most useful tools incorporated in SOFIA, which only requires a phased-allele matrix for all the markers in the map (in Figure 3F, light and dark grey color coding represents allele absence and presence, respectively). After the recombination blocks, a set of tiles in different colors represents the genes across the linkage groups in one of the maps (Figure 3G). Finally, links connecting similar markers between maps are draw in the interior of the figure (Figure 3H).

## Conclusions

SOFIA is an R package for running Perl native Circos within the R programming environment to display highly-dimensional genetic data. By integrating R and Circos, the package combines several advantages unavailable in other software packages. For example, SOFIA provides flexible formatting configuration and different plot styles for data exploration and for creating publication quality figures. It does not require users to have advanced computational abilities, but at the same time, it offers the possibility for experienced programmers and those familiar with native Circos to fully exploit the advantages of R to acquire complex figures. Finally, SOFIA fills a gap between the highly specialized but difficult to implement software Circos, and those such as ClicO FS, which are interactive and user friendly but also inflexible in terms of automatization.

## Availability and requirements

Project name: SOFIA: an R package for enhancing genetic visualization

Project home page: https://cran.r-project.org and https://cggl.horticulture.wisc.edu/software/

Operating system(s): Unix and Windows systems

Programming language: R

Other requirements: R > 2.0 and Circos

License: GPL-3

## Funding

This project was supported by USDA-ARS (project no. 3655-21220-001-00 provided to J.Z.); WI-DATCP (SCBG Project #14-002); Ocean Spray Cranberries, Inc.; Wisconsin Cranberry Growers Association; Cranberry Institute. BS was supported by the Frank B. Koller Cranberry Fellowship Fund for Graduate Students; GCP and LDG were supported by the Consejo Nacional de Ciencia y Tecnología (Mexico).

## Authors’ contributions

LDG, GCP, BS and JZ conceived the outline and main purpose of the software; LDG designed and implemented the software package; GCP participated in the software design and test. JZ and BS critically revised the manuscript. LDG and BS wrote the manuscript. All authors read and approved the final manuscript.

## Acknowledgements

JZ and BS wish to express their gratitude through 1 Cor 10:31. LDG wants to thank Martin Krzywinski for his advice and comments. We thank the anonymous reviewers who helped enhance the quality of this paper.

